# *Drosophila* pVALIUM10 TRiP RNAi lines cause undesired silencing of Gateway-based transgenes

**DOI:** 10.1101/2022.08.12.503771

**Authors:** Dimitrije Stanković, Gábor Csordás, Mirka Uhlirova

## Abstract

Post-transcriptional gene silencing using double-stranded RNA has revolutionized the field of functional genetics, allowing fast and easy disruption of gene function in various organisms. In *Drosophila*, many transgenic RNAi lines have been generated in large-scale efforts, including the *Drosophila* Transgenic RNAi Project (TRiP), to facilitate *in vivo* knockdown of virtually any *Drosophila* gene with spatial and temporal resolution. The available transgenic RNAi lines represent a fundamental resource for the fly community, providing an unprecedented opportunity to address a vast range of biological questions relevant to basic and biomedical research fields. However, caution should be applied regarding the efficiency and specificity of the RNAi approach. Here, we demonstrate that pVALIUM10-based RNAi lines, representing ~13% of the total TRiP collection (1,808 out of 13,410 TRiP pVALIUM-based RNAi lines), cause unintended off-target silencing of transgenes expressed from Gateway destination vectors generated via site-specific recombination. The silencing is mediated by targeting attB1 and attB2 sequences generated in the recombination reaction and included in the transcribed mRNA. Deleting these attB sites from the Gateway expression vector prevents silencing and restores expected transgene expression.

## INTRODUCTION

Double-stranded RNA (dsRNA)-mediated post-transcriptional gene silencing, also known as RNA interference (RNAi), represents an ancient anti-viral defense mechanism that has been harnessed as a powerful tool for reverse functional genetics in various cultured cells, plant and animal model systems (Kennerdell & Carthew, 1998; Lohmann *et al*, 1999; Wargelius *et al*, 1999; Wianny & Zernicka-Goetz, 2000; Zamore *et al*, 2000). The dsRNA silencing relies on a highly conserved RNase III enzyme called Dicer that recognizes and processes dsRNA into small 21-23-bp segments (siRNA). Upon loading into a multiprotein RNA-induced silencing complex (RISC), the guide strand facilitates pairing with complementary on-target RNAs, which are subsequently degraded by the action of the Argonaut proteins (Tomari & Zamore, 2005). In *Drosophila*, spatio-temporally controlled RNAi knockdown can be achieved by expressing dsRNA using transgenic binary expression systems, such as Gal4/UAS (Brand & Perrimon, 1993). The initial transgenic RNAi lines were generated using plasmids with ligated inverted repeats transcribed from one (Kennerdell & Carthew, 1998) or two opposite promoters (Giordano *et al*, 2002). However, these vectors were unstable and not well tolerated by the bacterial strains commonly used for cloning (Hagan & Warren, 1983; Piccin *et al*, 2001), a problem later solved by the introduction of an intron spacer between the repeats (Bao & Cagan, 2006; Lee & Carthew, 2003). This step-by-step progress led to the development of the pVALIUM (Vermilion-attB-LoxP-Intron-UAS-MCS), a universal vector system for transgenic RNAi library generation. The pVALIUM RNAi vectors contain an eye-color selection marker (*vermillion*), an attB site enabling phiC31-mediated site-directed integration into predefined genomic locations, *white* or *ftz* introns promoting the processing of dsRNA, and a cassette of two UAS pentamers, one of which is flanked by loxP sites to allow tuning of the dsRNA expression levels, as well as a multiple cloning site (MCS) (Ni *et al*, 2008).

The highly efficient pVALIUM10-based transgenic RNAi stocks produced within the *Drosophila* Transgenic RNAi Project (TRiP) at Harvard Medical School (HMS) (https://fgr.hms.harvard.edu) represent an invaluable resource for the *Drosophila* community (Perkins *et al*, 2015). At the moment of writing, 1,808 transgenic TRiP RNAi stocks based on the pVALIUM10 are available at the Bloomington *Drosophila* stock center (BDSC, https://bdsc.indiana.edu/index.html) (Perrimon *et al*, 2010). The pVALIUM10 vector carries *gypsy* insulators that markedly enhance knockdown efficiency, along with two inverted “attR1-selection-attR2” cassettes that facilitate cloning of dsRNA fragments via site-specific Gateway *in vitro* recombineering (Ni *et al*, 2009; Reece-Hoyes & Walhout, 2018). A desired single fragment, 400 - 600 bp in length and flanked by attL1 and attL2 recombination sites, can be easily moved from an entry vector into a pVALIUM10 destination vector via recombination between attL and attR sites catalyzed by LR clonase, producing a vector for UAS-driven expression of dsRNA (Ni *et al*., 2009).

The Gateway cloning technology is also routinely used to generate transgenic constructs for expression of epitope-tagged or untagged proteins, taking advantage of the *Drosophila* Gateway™ plasmid collection at the *Drosophila* Genomics Resource Center (https://dgrc.bio.indiana.edu/). Importantly, as with pVALIUM10 dsRNA plasmids, production of Gateway expression vectors relies on the recombination between attL and attR sites resulting in the formation of attB1 and attB2 sites.

Here, we demonstrate that the pVALIUM10-derived RNAi lines cause undesirable knockdown of transgenic reporters and overexpression lines generated with the help of the Gateway cloning technology. They do so by targeting the attB sites that are part of the transcribed mRNA. Deletion of the attB1 and attB2 sequences from the Gateway expression destination vector abrogates this effect and restores expected transgene expression levels.

## RESULTS

### Unexpected, off-target silencing of the RNAse H1::GFP reporter by multiple pVALIUM10-RNAi lines

*In vivo* RNAi-based genetic screens have proved very successful in identifying novel genes involved in developmental and pathologic processes in a variety of *Drosophila* somatic tissues when expressed using tissue-specific Gal4 driver lines (Graca *et al*, 2021; Lesch *et al*, 2010; Neely *et al*, 2010; Pletcher *et al*, 2019; Port *et al*, 2011; Rotelli *et al*, 2019; Rylee *et al*, 2022; Saj *et al*, 2010; Zeng *et al*, 2015; Zhou *et al*, 2019). Hence, we embarked on a candidate genetic screen aimed at identifying factors controlling genome stability. During the pre-screening of potential candidates we observed that expression of pVALIUM10-based *BuGZ^RNAi[TRiP.JF02830]^* RNAi line in the pouch region of the developing third instar wing discs, using the *nubbin-Gal4, UAS-mRFP (nub>mRFP*) driver line, efficiently suppressed the multi-epitope tagged BuGZ transgene (*BuGZ^FlyFos[v318366]^*) (Sarov *et al*, 2006) (Figure 1A, B, D). A similar reduction was observed using an independent *BuGZ^RNAi[KK104498]^* line from the Vienna *Drosophila* Resource Center (VDRC) KK RNAi library (Figure 1C, D). Encouraged by these results, both BuGZ RNAi lines were included in the candidate genetic screen that was based on expression of selected dsRNAs with the *nub>mRFP* driver. As readout for genome stability, we used the ubiquitously driven RNase H1::GFP that was generated by recombining the *RNase H1* coding sequence into the pUWG vector from the *Drosophila* Gateway™ plasmid collection. This resulted in the expression of a C-terminally GFP-tagged RNase H1 under the control of the poly-ubiquitin promoter (Figure 2A, B). The RNase H1 enzyme recognizes and resolves R-loops, DNA:RNA hybrids, that emerge during transcription and represent a threat to genome stability when unresolved (Crossley *et al*,2019; Santos-Pereira & Aguilera, 2015). Unexpectedly, we observed reduction of RNase H1::GFP signal when expressing *BuGZ^RNAi[TRiP.JF02830]^* but not *BuGZ^RNAi[KK104498]^* line (Figure 2C, D, H). Strikingly, similar downregulation of RNase H1::GFP was caused by pVALIUM10-based *myc^RNAi[TRiP.JF01761]^* but not *myc^RNAi[TRiP.HMS01538]^* or *myc^RNAi[GD2948]^* derived from pVALIUM20 and pMF3 vectors, respectively (Figures 1D and 2E-H). All three myc-RNAi lines have been shown effective by others (Aughey *et al*, 2016; Kim-Yip & Nystul, 2018; Ledru *et al*, 2022; Rust *et al*, 2018). These results indicated that the tested pVALIUM10 TRiP RNAi lines but not pVALIUM20, KK or GD lines may cause undesirable silencing of the RNase H1::GFP transgene. We further speculated that dsRNA expressed from pVALIUM10 contains segments unrelated to the target gene-specific sequences that trigger this unexpected off-target effect.

**Figure 1.**
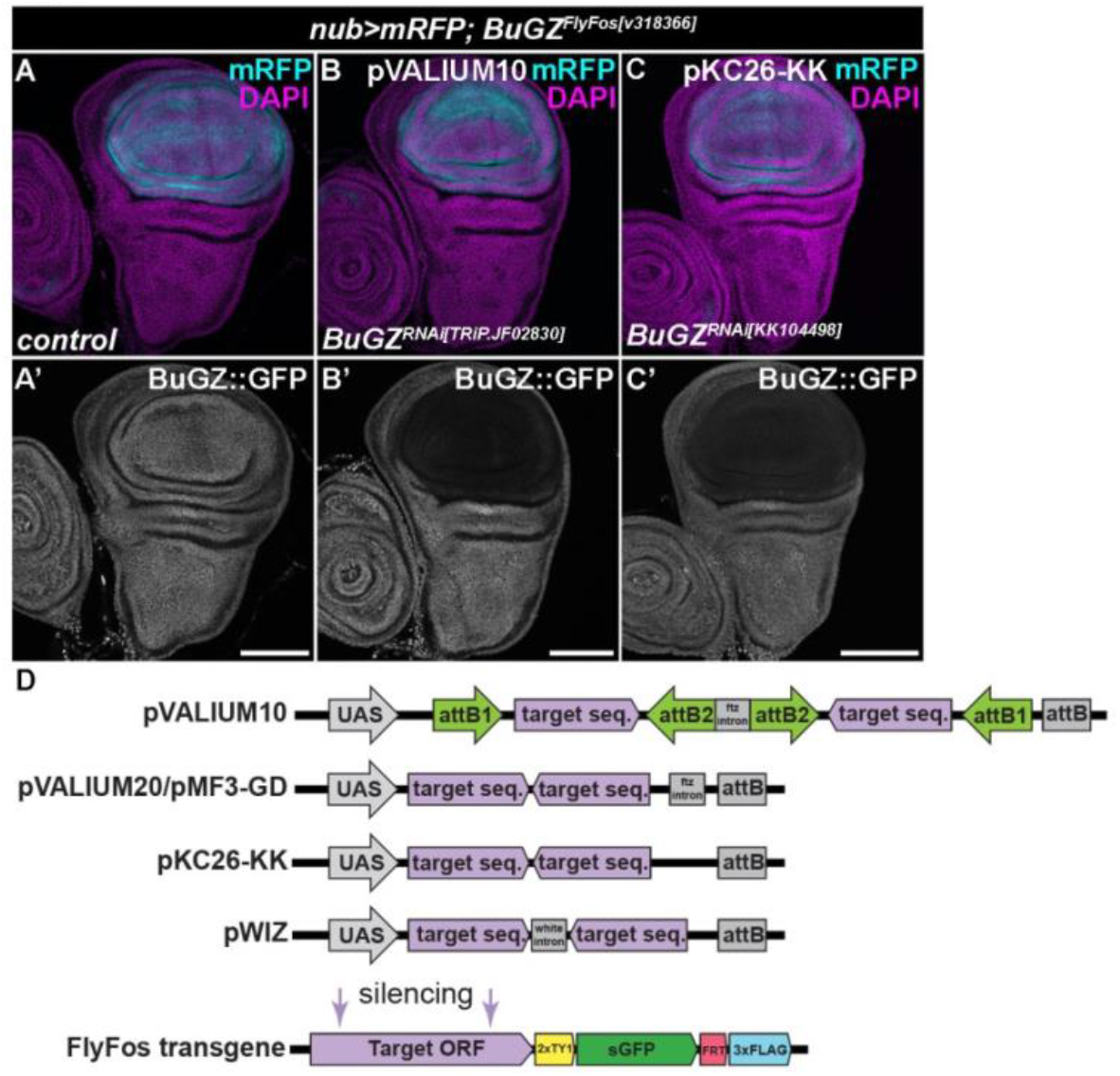
pVALIUM10 and KK RNAi lines cause efficient silencing. **(A-C)** Expression of pVALIUM10 *BuGZ^RNAi[TRIP.JF02830]^* (B, B’) and KK-based *BuGZ^RNAi[KK104498]^* transgenic RNAi lines (C, C’) using the *nubbin-Gal4, UAS-myr-mRFP* driver (*nub>mRFP*, light blue) causes a marked reduction of a BuGZ^FlyFos[v318366]^ transgenic protein in the pouch region of wing discs (WDs) relative to the hinge and notum, and control WD (A, A’). Micrographs show projections of multiple confocal sections of WDs dissected from third instar larvae 7 days AEL. The multiple-tagged BuGZ^FlyFos [v318366]^ protein was visualized by immunostaining with an anti-GFP antibody (white). Nuclei were counterstained with DAPI (magenta). Scale bars: 100 μm. **(D)** Schematic representation of various vectors used for cloning and overexpression of dsRNA. In the absence of the target-specific antibody, the RNAi efficacy can be assessed with the help of FlyFos transgenes expressing multiple-tagged proteins under the control of the endogenous regulatory sequences.

**Figure 2.**
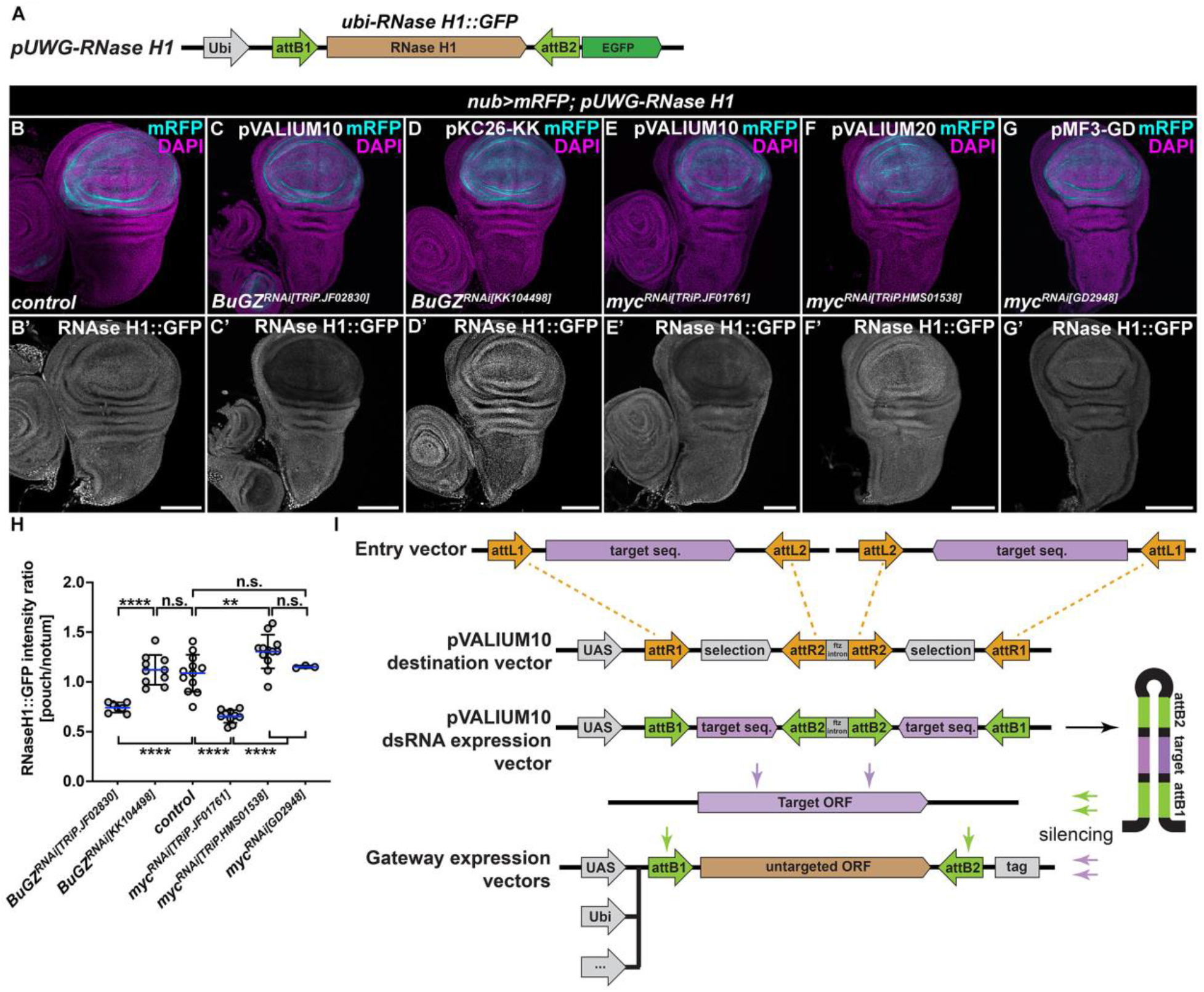
pVALIUM10-based lines cause off-target silencing of the RNAse H1::GFP transgenic reporter. **(A)** Schematic representation of the transgenic *pUWG-RNase H1* reporter construct expressing the C-terminally-tagged RNase H1::GFP protein from the ubiquitous (*ubi*) promoter. The *RNase H1* coding sequence is flanked by attB1 and attB2 sites generated via Gateway-mediated recombination catalyzed by the LR clonase. **(B-G)** Immunostaining with a GFP-specific antibody revealed a marked reduction of the RNase H1::GFP levels in the nubbin domain (mRFP, light blue), where pVALIUM10-based *BuGZ^RNAi[TRIP.JF02830]^* (C) and *myc^RNAi[TRiP.JF01761]^* (E) RNAi lines were expressed using the *nubbin-Gal4, UAS-myr-mRFP* driver (*nub>mRFP*). No such downregulation of the RNase H1::GFP signal was observed in control WD (B) and following expression of KK-based *BuGZ^RNAi[KK104498]^* (D), pVALIUM20-based *myc^RNAi[TRiP.HMS01538]^* (F) or GD-based *myc^RNAi[GD2948]^* (G). Micrographs show projections of multiple confocal sections of WDs dissected from third instar larvae 7 days AEL. Nuclei were counterstained with DAPI (magenta). Scale bars: 100 μm. **(H)** Quantification of the RNase H1::GFP signal depicted as relative intensity ratio between the pouch and notum region. Statistical significance was determined by one-way ANOVA with Tukey’s multiple comparison test; n ≥ 3; **p<0.01, ****p<0.0001, n.s. non-significant. **(I)** Generation of pVALIUM10 RNAi lines relied on a LR clonase-mediated recombination of dsRNA fragment from an Entry vector to the pVALIUM10 destination vector (Ni *et al*., 2009). The dsRNA hairpin produced from the pVALIUM10 dsRNA expression vector is the source of siRNA species targeting the transcript of interest (target ORF) but also attB1 and attB2 sites present in any of the Gateway expression vectors generated via the same recombination mechanisms.

### pVALIUM10 RNAi and Gateway expression plasmids share attB recombination sites

To identify the potential target for dsRNA-mediated knockdown of the RNAse H1::GFP transgene by pVALIUM10 TRiP RNAi, we compared the plasmid sequences. We found that attB1 and attB2 sites are shared between the Gateway-based expression vectors, including pUWG, and pVALIUM10 TRiP RNAi plasmids. AttB1 and attB2 sites are 25 bp in length and generated by LR clonase-mediated recombination between attL and attR sequences. Given the repetition of attB1 and attB2 sites, flanking the passenger and guide target sequence fragments in pVALIUM10 TRiP RNAis, these sequences are transcribed and could contribute to the dsRNA stem-loop substrate for Dicer. Based on these results we hypothesized that dsRNAs generated from attB1 and attB2 could target, and thus silence, Gateway-based transgenes via attB sites (Figure 2I).

### pVALIUM10 RNAi lines silence Gateway transgenes via attB sites

To test our hypothesis, we cloned the *mCherry* coding sequence into the pUWG Gateway expression vector (*pUWG-mCherry*) allowing ubiquitous expression of mCherry transcript flanked by the attB1 and attB2 sites. The pUWG*-mCherry* vector then served as a template for overlap PCR to generate the pUWG^ΔattB^-*mCherry* construct that lacks attB sites (Figure 3A). Both plasmids were used to establish transgenic fly lines through a standard *Drosophila* germline transformation method (Rubin & Spradling, 1982). The susceptibility or resistance of *ubi-mCherry (pUWG-mCherry*) and *ubi^ΔattB^-mCherry* (pUWG^Δatt*B*^-*mCherry*) transcripts to RNAi-mediated silencing was then tested by driving various RNAi lines under the control of the *nub-Gal4* driver. Immunostaining of the third instar wing imaginal discs using an anti-RFP antibody revealed an identical pattern of ubi-mCherry and ubi^ΔattB^-mCherry transgenes in the control (*nub>*) background (Figure 3B, C). In contrast, levels of ubi-mCherry but not ubi^ΔattB^-mCherry transgenic protein were markedly reduced in the wing pouch region relative to the rest of the tissue following overexpression of *BuGZ* and *myc* pVALIUM10 TRiP RNAi lines (Figure 3D, E, H, I, J, M) while no significant changes were observed when *BuGZ* and *myc* were knocked down using KK and pVALIUM20 RNAi lines, respectively (Figure 3F, G, H, K, L, M). Importantly, the attB site-dependent silencing of ubi-mCherry was recapitulated using an additional set of pVALIUM10 TRiP RNAi lines targeting *Atf3, Lip4, yki, DH44, ftz-f1, mago, brat*, and *Actβ* genes (Supplementary Figure S1).

**Figure 3.**
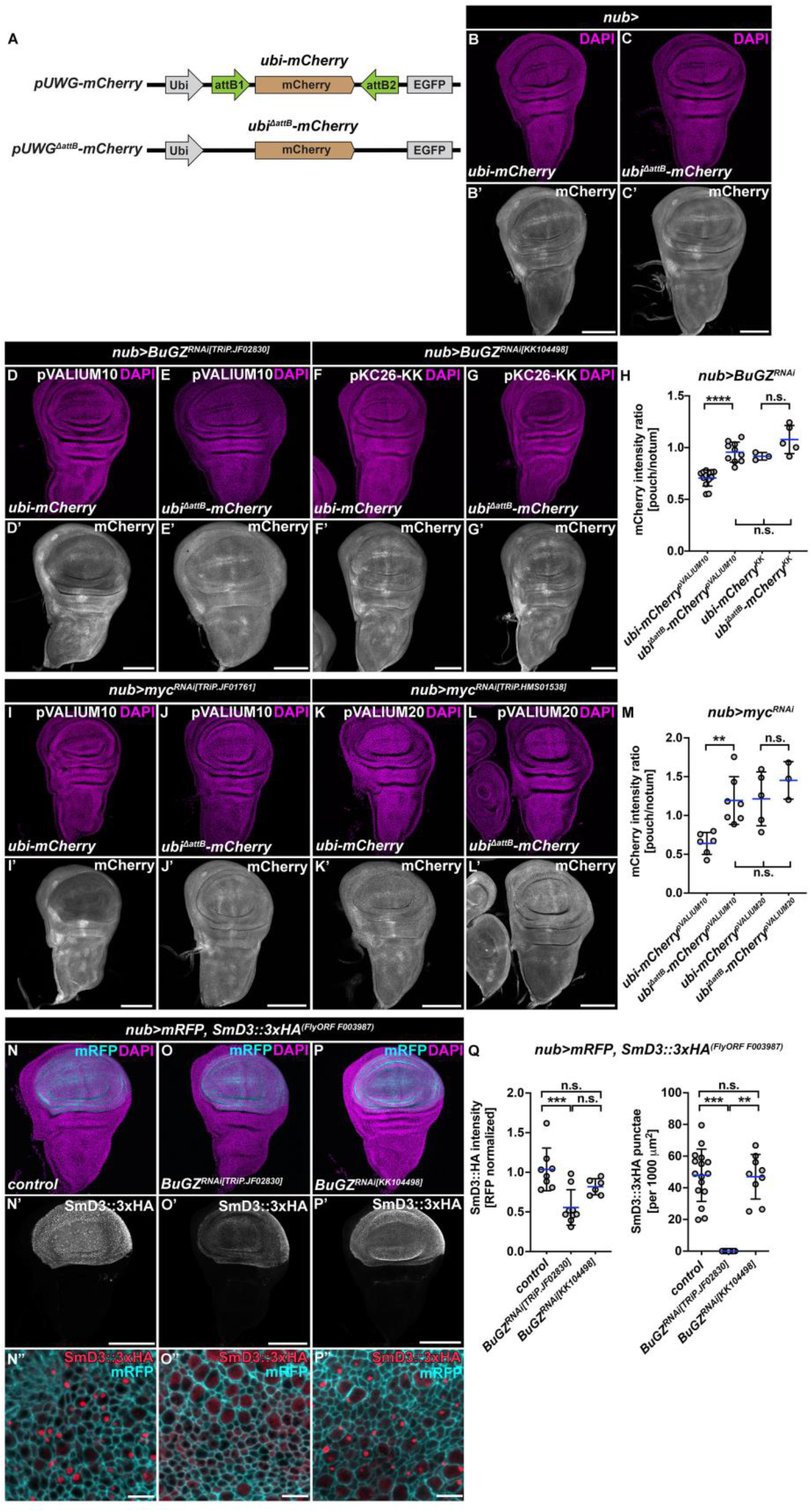
pVALIUM10 RNAi lines silence Gateway transgenes via attB1 and attB2 sites. **(A)** Schematic of the transgenic *pUWG-mCherry* and mutant *pUWG^ΔattB^-mCherry* reporter constructs. The mCherry is expressed from the poly-ubiquitin (*ubi*) promoter. **(B, C)***ubi-mCherry* and mutant *ubi^ΔattB^-mCherry* reporters showed similar expression pattern in the wing imaginal discs. **(D-M)***ubi-mCherry* was efficiently downregulated by pVALIUM10 *BuGZ^RNAi[TRIP.JF02830]^* and *myc^RNAi[TRiP.JF01761]^* expressed under the control of the *nubbin-Gal4* driver (*nub>*) relative to the rest of the WD (D, I, H, M). In contrast, absence of attB1 and attB2 sequences rendered the *ubi^ΔattB^-mCherry* reporter resistant to pVALIUM10 RNAi-mediated silencing (E, H, J, M). Neither KK- (*BuGZ^RNAi[KK104498]^*) nor pVALIUM20-based (*myc^RNAi[TRiP.HMS01538]^*) RNAi lines affected mCherry levels expressed from the unmodified or mutant reporter constructs (F-H, K-M). Micrographs show projections of multiple confocal sections of WDs dissected from third instar larvae 7 days AEL that were immunostained with an anti-mRFP antibody (white). Nuclei were counterstained with DAPI (magenta). Scale bars: 100 μm. Quantification of mCherry signal depicted as relative intensity ratio between the pouch and notum region (H, M). Superscript text in each group denotes the type of RNAi used. Statistical significance was determined by one-way ANOVA with Tukey’s multiple comparison test; n ≥ 3; **p<0.01, ****p<0.0001, n.s. non-significant. **(N-Q)** Expression of pVALIUM10 *BuGZ^RNAi[TRIP.JF02830]^* (O) in the wing pouch using *nubbin-Gal4, UAS-mRFP* driver (*nub>mRFP*) reduces SmD3::3xHA levels compared to control (N) and KK-based *BuGZ^RNAi[KK104498]^* (P). Note foci of SmD3::3xHA protein in control (N”) and *BuGZ^RNAi[KK104498]^* cells (P”) cells that are not present in *BuGZ^RNAi[TRIP.JF02830]^* expressing cells (O”). Micrographs show projections of multiple or single (N”-P”) confocal sections of WDs dissected from third instar larvae 7 days AEL that were immunostained with an anti-HA antibody (white, N’-P’; red, N”-P”). Nuclei were counterstained with DAPI (magenta). Scale bars: 100 μm or 5 μm (N”-P”). Quantification of the SmD3::3xHA signal intensity (Q, left) and SmD3::3xHA-positive punctae (Q, right) in the pouch region. Statistical significance was determined by one-way ANOVA with Tukey’s multiple comparison test and Kruskall-Wallis test, respectively; n ≥ 6; **p<0.01, ***p<0.001, n.s. non-significant.

Finally, we tested whether pVALIUM10 TRiP RNAi lines may also interfere with UAS-inducible attB-containing transcripts, such as transgenic fly open reading frames (ORFs) expressed from the Gateway destination vector pGW-HA.attB (Bischof *et al*, 2014). To this end, we assessed levels of the HA-tagged SmD3 FlyORF protein in the presence of pVALIUM10 or KK BuGZ-RNAi line. In contrast to an enrichment of SmD3::3xHA protein and positive punctae in the wing pouch cells of control (*nub>*) and *BuGZ^RNAi[KK104498]^*expressing wing discs, the overall levels and the number of positive punctae were noticeably reduced in *nub>BuGZ^RNAi[TRiP.JF02830]^* cells (Figure 3N-Q). These results suggest that the dsRNA hairpin produced from pVALIUM10 RNAi vectors generates attB1- and attB2-siRNAs, which guide destruction of transcripts containing such target sites. This off-target silencing via attB sites is specific to RNAi lines based on pVALIUM10. Absence of attB1 and attB2 sites renders the Gateway-derived transcript resistant to pVALIUM10 TRiP RNAi.

## DISCUSSION

The use of transgenic RNAi to achieve developmental stage- and tissue-specific knockdown of candidate genes and to perform genome-wide screens in *Drosophila* has become commonplace in the past two decades (Blake *et al*, 2017; Dietzl *et al*, 2007; Hu *et al*, 2017; Port *et al*., 2011). Both our knowledge on the RNAi mechanism, and the tools to generate new types of RNAi constructs efficiently have considerably evolved in recent years. As a result, several independent transgenic RNAi stock collections were established, including Vienna *Drosophila* stock center (VDRC, 23,411 RNAi lines, https://stockcenter.vdrc.at/control/main), Bloomington *Drosophila* Stock Center (BDSC, 13,698 RNAi lines, https://bdsc.indiana.edu/), and the National Institute of Genetics (NIG, 12,365 RNAi lines, https://shigen.nig.ac.jp/fly/nigfly/) at the time of writing. Yet, in spite of being a broadly used technique, transgenic manipulation of gene expression can still have unforeseen consequences. A large portion of the RNAi lines from the VDRC KK collection were found to produce phenotypes due to a secondary insertion and consequent ectopic expression of the *tiptop* gene (Green *et al*,2014; Vissers *et al*, 2016). More recently, van der Graaf and colleagues reported that transgenes inserted in the commonly used *attP40* landing site on the second chromosome can decrease the expression of Msp300, a member of the LINC complex, and hence lead to nuclear size, shape and positioning phenotypes (van der Graaf *et al*,2022).

Here, we demonstrate that the pVALIUM10-based RNAi lines cause off-target silencing of ORFs expressed from Gateway expression vectors or potentially any transcripts containing attB1 and attB2 sites. We show that the removal of these sites abrogates the undesirable knockdown. Given the popularity of the Gateway expression vectors to generate transgenic protein-reporters and tagged overexpression constructs, our results suggest that simultaneous use of pVALIUM10 TRiP RNAi may lead to experimental artefacts unrelated to the investigated biological process.

Of note, in the original publication of pVALIUM10 vectors, the authors tested for potential off-target effects by comparing the hairpin sequence against the *Drosophila* genome (Ni *et al*., 2009), which does not contain endogenous 21nt stretches equivalent to the attB1 and attB2 sequences. However, as the transgenic toolkit in *Drosophila* is ever expanding, each additional transgene has the possibility to produce a synthetic effect with the existing RNAi lines. Hence, we suggest that the researchers validate the type of TRiP RNAi line or the expression vector sequence before commencing their experiments. As an example, we do not recommend combining RNAi based on pVALIUM10 with the 3xHA-tagged overexpression lines from FlyORF (https://flyorf.ch) (Bischof *et al*., 2014) or the fly-FUCCI (Fluorescent Ubiquitination-based Cell Cycle Indicator) lines, which allow real-time visualization of the cell cycle dynamics by the expression two fluorescently tagged degrons (Zielke et al., 2014). In both cases, the transgenes were generated via Gateway reaction and contains attB sites flanking the inserted ORF (Bischof *et al*., 2014; Zielke *et al*, 2014), which can lead to off-target silencing of the transcript. Furthermore, this finding reinforces the idea that RNAi experiments should adhere to the standards established in the past decades, including investigating possible off-target effects to eliminate knockdown of the unintended gene, using multiple independent RNAi lines to recapitulate phenotypes, and confirming the knockdown of the targeted gene products (Echeverri *et al*, 2006; Green *et al*., 2014; Kaya-Çopur & Schnorrer, 2016).

## MATERIALS AND METHODS

### Fly stocks and Husbandry

The following *Drosophila* strains were used: *w^1118^* (BDSC; RRID: BDSC_3605), *nubbin-Gal4, UAS-myr-mRFP (nub>mRFP), BuGZ^FlyFos[v318366]^* (VDRC), *BuGZ^RNAi[TRiP.JF02830]^* (BDSC; RRID:BDSC_27996), *BuGZ^RNAi[KK104498]^* (VDRC), *Lip4^RNAi[TRiP.HMO5136]^* (BDSC; RRID:BDSC_28925), *myc^RNAi[TRiP.JF01761]^* (BDSC; RRID:BDSC_25783), *myc^RNAi[TRiP.HMS01538]^* (BDSC; RRID:BDSC_36123), *myc^RNAi[GD2948]^* (VDRC), *Actß^RNAi[TRiP.JF03276]^* (BDSC; RRID:BDSC_29597), *ftz-f1^RNAi[TRiP.JF02738]^* (BDSC; RRID:BDSC_27659), *yki^RNAi[TriP.JF03119]^* (BDSC; RRID:BDSC_31965), *Atf3^RNAi[TRiP.JF02303]^* (BDSC; RRID:BDSC_26741), *DH44^RNAi[TRiP.JF01822]^* (BDSC; RRID:BDSC_25804), *brat^RNAi[TRiP.HM05078]^* (BDSC; RRID:BDSC_28590), *mago^RNAi[TRiP.HM05142]^* (BDSC; RRID:BDSC_28931), *SmD3::3xHA^[FlyORF F003987]^* (FlyORF), *nub>SmD3::3xHA^[FlyORF F003987]^* (Erkelenz *et al*, 2021), *ubi-RNase H1::GFP, ubi-mCherry, ubi^ΔattB^-mCherry* (this study). All *Drosophila* stocks are listed in Supplementary Table S1. All crosses were set up and maintained at 25 °C, unless specified otherwise, on a diet consisting of 0.8% agar, 8% cornmeal, 1% soymeal, 1.8% dry yeast, 8% malt extract, and 2.2% sugar-beet syrup, which was supplemented with 0.625% propionic acid and 0.15% Nipagin. *w^1118^* line was used as control.

### Generation of plasmids and transgenic lines

The open reading frame of *rnh1* (FBgn0023171) was amplified from *Drosophila melanogaster* cDNA (*w^1118^*)using primers harboring NotI and SalI restriction sites. The stop codon was excluded to allow C-terminal tagging of the protein. The ORF was cloned into the Gateway pENTR4 Dual Selection Vector (Thermo Fisher Scientific; #A10465). The ORF was then recombined into the pUWG vector (DGRC; #1284) using the LR Clonase II enzyme mix according to manufacturer’s instructions (Thermo Fisher Scientific; #11791020). The coding sequence of *mCherry* (GenBank: AY678264.1) was amplified with specific primers, cloned into the pENTR 4 Dual Selection vector (Invitrogen, Cat. Nr. #A10465) including a stop codon and subsequently recombined into the pUWG vector (*pUWG-mCherry*). Overlap PCR strategy was employed to generate the pUWG^ΔattB^*-mCherry* construct lacking attB sites resulting from the LR recombination. Three separate PCR reactions were performed to produce fragments for assembly using the original *pUWG-mCherry* plasmid as a template. The forward primer of the first primer pair contained a XhoI restriction site, while the reverse primer contained a XbaI restriction site. Partial complementarity of the reverse and forward primers used to synthesize the downstream fragment enabled the exclusion of attB sites in the final product. The final PCR product was ligated into the pUWG*-mCherry* plasmid digested with XbaI and XhoI. Transgenic fly lines with random insertions of the constructs were generated by standard P-element-mediated transformation using *w^1118^ Drosophila* strain. All primers and plasmids are listed in Supplementary Table S2.

### Tissue dissection and immunostaining

Wing imaginal discs dissected from third instar *Drosophila* larvae (7 days after egg laying (AEL)) were fixed for 25 minutes with 4% paraformaldehyde in PBS containing 0.1% Triton X (PBS-T) at room temperature. Fixed tissues were washed 3 times with PBS-T. Primary antibodies were diluted in blocking buffer (PBS-T with 0.3% BSA) and tissues were stained overnight at 4 °C. The following primary antibodies were used: rabbit anti-mRFP (1:500, MBL International Cat# PM005, RRID:AB_591279), goat anti-GFP (1:500, Abcam Cat# ab6673, RRID:AB_305643), and rabbit-anti HA (1:1000, Abcam Cat# ab9110, RRID:AB_307019). After washing, the samples were incubated with the corresponding Alexa Fluor 488 (1:2000, Thermo Fisher Scientific Cat# A-11034, RRID:AB_2576217), Cy2 (1:2000, Jackson ImmunoResearch Labs Cat# 705-225-147, RRID:AB_2307341) or Cy3 (Jackson ImmunoResearch Labs Cat# 711-165-152, RRID:AB_2307443) conjugated secondary antibodies overnight at 4 °C, washed and counterstained with DAPI (1:1000 dilution of 5 mg/ml stock, 6335.1, Carl Roth GmbH) to visualize nuclei. Tissues were mounted on glass slides in Dabco-Mowiol 4-88 (Sigma-Aldrich Cat# D2522 and Cat# 81381).

### Image acquisition and processing

Confocal images and stacks were acquired with Olympus FV1000 confocal microscope equipped with 20X UPlan S-Apo (NA 0.85), 40X UPlan FL (NA 1.30) and 60X UPlanApo (NA1.35) objectives. Maximum Z-projections were generated from consecutive sections taken at 1.4 μm steps using Fluoview 1000 Software (Olympus) (RRID: SCR_014215) and Fiji (https://fiji.sc/) (RRID:SCR_002285). Final image processing, including panel assembly, brightness and contrast adjustments were performed in Adobe Photoshop CC (Adobe Systems, Inc.) (RRID:SCR_014199).

### Quantification of mCherry, RNase H1::GFP, and SmD3::3xHA signal

Fluorescent signal intensity was quantified with Fiji (https://fiji.sc/) (RRID:SCR_002285). For mCherry and RNase H1::GFP, a square selection was created in the middle of the pouch region, and the,, Mean intensity” was measured, then the same selection was moved to the middle of the notum and the measurement was repeated. The two values were divided to produce a pouch/notum intensity ratio for each wing disc. For SmD3::3xHA, the,, Mean intensities” of SmD3::3xHA and mRFP signals in the pouch region were measured and divided to produce a normalized value for each wing disc. Zoom images of the nubbin domain of independent WDs were used to count the number of SmD3::3xHA-positive punctae per 1000 μm^2^ area. Statistical significance was determined by one-way ANOVA with Tukey’s multiple comparison test (signal intensity) or Kruskall-Wallis test (puncta) in GraphPad Prism (RRID:SCR_002798).

## ACKNOWLEDGEMENTS

We thank Norbert Perrimon for discussions and comments on the manuscript. We thank the Bloomington *Drosophila* Stock Center (BDSC, Bloomington, IN, USA), the Vienna *Drosophila* Resource Center (VDRC, Vienna, Austria), *Drosophila* Genomic Resource Center (DGRC, supported by NIH grant 2P40OD010949, Bloomington, IN, USA), the Zurich ORFeome Project (FlyORF, Zurich, Switzerland) for fly stocks and plasmids. We are grateful to Tina Bresser for generation of transgenic lines and Nils Teuscher for fly stock maintenance and technical assistance.

## Notes

### Competing Interest Statement

The authors have declared no competing interest.

### Summary of Updates

The absolute number of pVALIUM10 lines was included in the manuscript abstract. The percentages have been corrected. The total number of RNAi lines available from various stock centers mentioned in the Discussion was updated/corrected. Figure 2 revised (quantification has been included, now panel H) Figure 3 revised (labeling in panels H and M was corrected and new quantification is now included as panel Q)

## REFERENCES

Aughey GN, Grice SJ, Liu J-L (2016) The Interplay between Myc and CTP Synthase in Drosophila. PLOS Genetics 12: e1005867

Bao S, Cagan R (2006) Fast cloning inverted repeats for RNA interference. RNA 12: 2020–2024

Bischof J, Sheils EM, Björklund M, Basler K (2014) Generation of a transgenic ORFeome library in Drosophila. Nature protocols 9: 1607–1620

Blake AJ, Finger DS, Hardy VL, Ables ET (2017) RNAi-Based Techniques for the Analysis of Gene Function in Drosophila Germline Stem Cells. Methods in molecular biology (Clifton, NJ) 1622: 161–184

Brand AH, Perrimon N (1993) Targeted gene expression as a means of altering cell fates and generating dominant phenotypes. Development 118: 401–415

Crossley MP, Bocek M, Cimprich KA (2019) R-Loops as Cellular Regulators and Genomic Threats. Mol Cell 73: 398–411

Dietzl G, Chen D, Schnorrer F, Su KC, Barinova Y, Fellner M, Gasser B, Kinsey K, Oppel S, Scheiblauer S et al (2007) A genome-wide transgenic RNAi library for conditional gene inactivation in Drosophila. Nature 448: 151–156

Echeverri CJ, Beachy PA, Baum B, Boutros M, Buchholz F, Chanda SK, Downward J, Ellenberg J, Fraser AG, Hacohen N et al (2006) Minimizing the risk of reporting false positives in large-scale RNAi screens. Nature Methods 3: 777–779

Erkelenz S, Stanković D, Mundorf J, Bresser T, Claudius A-K, Boehm V, Gehring NH, Uhlirova M (2021) Ecd promotes U5 snRNP maturation and Prp8 stability. Nucleic Acids Research

Giordano E, Rendina R, Peluso I, Furia M (2002) RNAi triggered by symmetrically transcribed transgenes in Drosophila melanogaster. Genetics 160: 637–648

Graca FA, Sheffield N, Puppa M, Finkelstein D, Hunt LC, Demontis F (2021) A large-scale transgenic RNAi screen identifies transcription factors that modulate myofiber size in Drosophila. PLOS Genetics 17: e1009926

Green EW, Fedele G, Giorgini F, Kyriacou CP (2014) A Drosophila RNAi collection is subject to dominant phenotypic effects. Nat Methods 11: 222–223

Hagan CE, Warren GJ (1983) Viability of palindromic DNA is restored by deletions occurring at low but variable frequency in plasmids of Escherichia coli. Gene 24: 317–326

Hu Y, Comjean A, Roesel C, Vinayagam A, Flockhart I, Zirin J, Perkins L, Perrimon N, Mohr SE (2017) FlyRNAi.org-the database of the Drosophila RNAi screening center and transgenic RNAi project: 2017 update. Nucleic Acids Res 45: D672–D678

Kaya-Çopur A, Schnorrer F (2016) A Guide to Genome-Wide In Vivo RNAi Applications in Drosophila. Methods Mol Biol 1478: 117–143

Kennerdell JR, Carthew RW (1998) Use of dsRNA-mediated genetic interference to demonstrate that frizzled and frizzled 2 act in the wingless pathway. Cell 95: 1017–1026

Kim-Yip RP, Nystul TG (2018) Wingless promotes EGFR signaling in follicle stem cells to maintain self-renewal. Development 145: dev168716

Ledru M, Clark CA, Brown J, Verghese S, Ferrara S, Goodspeed A, Su TT (2022) Differential gene expression analysis identified determinants of cell fate plasticity during radiation-induced regeneration in Drosophila. PLoS Genet 18: e1009989

Lee YS, Carthew RW (2003) Making a better RNAi vector for Drosophila: use of intron spacers. Methods 30: 322–329

Lesch C, Jo J, Wu Y, Fish GS, Galko MJ (2010) A targeted UAS-RNAi screen in Drosophila larvae identifies wound closure genes regulating distinct cellular processes. Genetics 186: 943–957

Lohmann JU, Endl I, Bosch TC (1999) Silencing of developmental genes in Hydra. Dev Biol 214: 211–214

Neely GG, Kuba K, Cammarato A, Isobe K, Amann S, Zhang L, Murata M, Elmen L, Gupta V, Arora S et al (2010) A global in vivo Drosophila RNAi screen identifies NOT3 as a conserved regulator of heart function. Cell 141: 142–153

Ni JQ, Liu LP, Binari R, Hardy R, Shim HS, Cavallaro A, Booker M, Pfeiffer BD, Markstein M, Wang H et al (2009) A Drosophila resource of transgenic RNAi lines for neurogenetics. Genetics 182: 1089–1100

Ni JQ, Markstein M, Binari R, Pfeiffer B, Liu LP, Villalta C, Booker M, Perkins L, Perrimon N (2008) Vector and parameters for targeted transgenic RNA interference in Drosophila melanogaster. Nat Methods 5: 49–51

Perkins LA, Holderbaum L, Tao R, Hu Y, Sopko R, McCall K, Yang-Zhou D, Flockhart I, Binari R, Shim HS et al (2015) The Transgenic RNAi Project at Harvard Medical School: Resources and Validation. Genetics 201: 843–852

Perrimon N, Ni JQ, Perkins L (2010) In vivo RNAi: today and tomorrow. Cold Spring Harb Perspect Biol 2: a003640

Piccin A, Salameh A, Benna C, Sandrelli F, Mazzotta G, Zordan M, Rosato E, Kyriacou CP, Costa R (2001) Efficient and heritable functional knock-out of an adult phenotype in Drosophila using a GAL4-driven hairpin RNA incorporating a heterologous spacer. Nucleic Acids Research 29

Pletcher RC, Hardman SL, Intagliata SF, Lawson RL, Page A, Tennessen JM (2019) A Genetic Screen Using the Drosophila melanogaster TRiP RNAi Collection To Identify Metabolic Enzymes Required for Eye Development. G3 (Bethesda) 9: 2061–2070

Port F, Hausmann G, Basler K (2011) A genome-wide RNA interference screen uncovers two p24 proteins as regulators of Wingless secretion. EMBO Rep 12: 1144–1152

Reece-Hoyes JS, Walhout AJM (2018) Gateway Recombinational Cloning. Cold Spring Harb Protoc 2018: pdb.top094912

Rotelli MD, Bolling AM, Killion AW, Weinberg AJ, Dixon MJ, Calvi BR (2019) An RNAi Screen for Genes Required for Growth of Drosophila Wing Tissue. G3 Genes|Genomes|Genetics 9: 3087–3100

Rubin GM, Spradling AC (1982) Genetic transformation of Drosophila with transposable element vectors. Science 218: 348–353

Rust K, Tiwari MD, Mishra VK, Grawe F, Wodarz A (2018) Myc and the Tip60 chromatin remodeling complex control neuroblast maintenance and polarity in Drosophila. EMBO J 37

Rylee J, Mahato S, Aldrich J, Bergh E, Sizemore B, Feder LE, Grega S, Helms K, Maar M, Britt SG et al (2022) A TRiP RNAi screen to identify molecules necessary for Drosophila photoreceptor differentiation. G3 Genes|Genomes|Genetics: jkac257

Saj A, Arziman Z, Stempfle D, van Belle W, Sauder U, Horn T, Dürrenberger M, Paro R, Boutros M, Merdes G (2010) A Combined Ex Vivo and In Vivo RNAi Screen for Notch Regulators in Drosophila Reveals an Extensive Notch Interaction Network. Developmental Cell 18: 862–876

Santos-Pereira JM, Aguilera A (2015) R loops: new modulators of genome dynamics and function. Nat Rev Genet 16: 583–597

Sarov M, Schneider S, Pozniakovski A, Roguev A, Ernst S, Zhang Y, Hyman AA, Stewart AF (2006) A recombineering pipeline for functional genomics applied to Caenorhabditis elegans. Nat Methods 3: 839–844

Tomari Y, Zamore PD (2005) Perspective: machines for RNAi. Genes Dev 19: 517–529

van der Graaf K, Srivastav S, Singh P, McNew JA, Stern M (2022) The Drosophila attP40 docking site and derivatives are insertion mutations of MSP300. bioRxiv: 2022.2005.2014.491875

Vissers JH, Manning SA, Kulkarni A, Harvey KF (2016) A Drosophila RNAi library modulates Hippo pathway-dependent tissue growth. Nat Commun 7: 10368

Wargelius A, Ellingsen S, Fjose A (1999) Double-stranded RNA induces specific developmental defects in zebrafish embryos. Biochem Biophys Res Commun 263: 156–161

Wianny F, Zernicka-Goetz M (2000) Specific interference with gene function by double-stranded RNA in early mouse development. Nat Cell Biol 2: 70–75

Zamore PD, Tuschl T, Sharp PA, Bartel DP (2000) RNAi: double-stranded RNA directs the ATP-dependent cleavage of mRNA at 21 to 23 nucleotide intervals. Cell 101: 25–33

Zeng X, Han L, Singh SR, Liu H, Neumuller RA, Yan D, Hu Y, Liu Y, Liu W, Lin X et al (2015) Genome-wide RNAi screen identifies networks involved in intestinal stem cell regulation in Drosophila. Cell Rep 10: 1226–1238

Zhou J, Xu L, Duan X, Liu W, Zhao X, Wang X, Shang W, Fang X, Yang H, Jia L et al (2019) Large-scale RNAi screen identified Dhpr as a regulator of mitochondrial morphology and tissue homeostasis. Science Advances 5: eaax0365

Zielke N, Korzelius J, van Straaten M, Bender K, Schuhknecht GFP, Dutta D, Xiang J, Edgar BA (2014) Fly-FUCCI: A versatile tool for studying cell proliferation in complex tissues. Cell Rep 7: 588–598

