## Supplementary Figure S1 Tables S1_S2 for "*Drosophila* pVALIUM10 TRiP RNAi lines cause undesired silencing of Gateway-based transgenes"

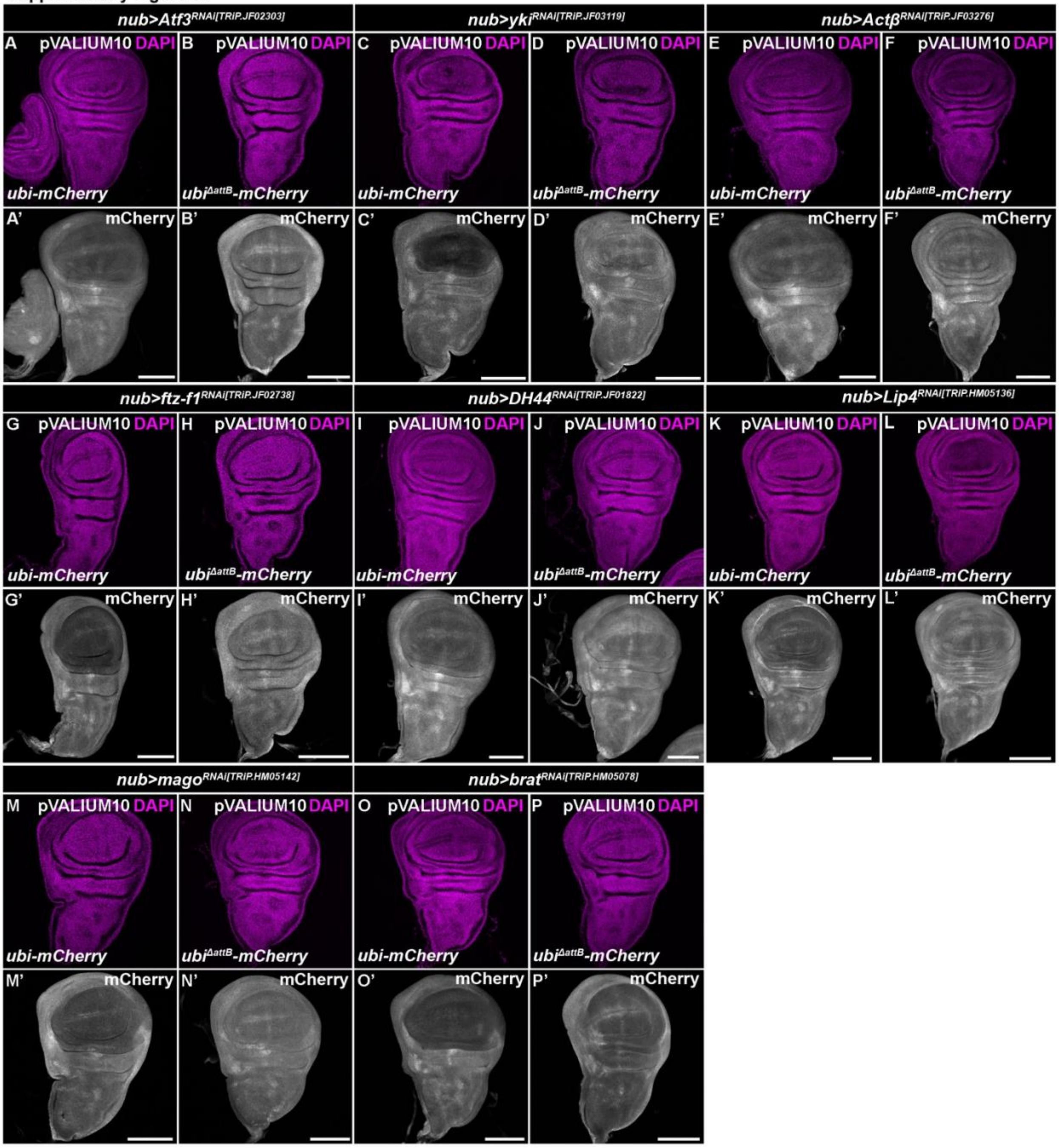


**Supplementary Figure S1. pVALIUM10 RNAi lines silence Gateway transgenes via attB1 and attB2 sites.**

**(A-P)** Wild-type *ubi-mCherry* but not the *ubi*$^{\Delta attB}$*-mCherry* reporter lacking attB1 and attB2 sequences was efficiently downregulated in the pouch region by pVALIUM10 RNAi lines targeting *Atf3* (A, B), *yki*, (C, D), *Act*$\beta$ (E, F), *ftz-f1* (G, H), *DH44* (I, J), *Lip4* (K, L), *mago* (M, N), *brat* (O, P) transcripts expressed under the control of the *nubbin-Gal4* driver (*nub*>) relative to the rest of the WD. Micrographs show projections of multiple confocal sections of WDs dissected from third instar larvae 7 days AEL that were immunostained with an anti-mRFP antibody (white). Nuclei were counterstained with DAPI (magenta). Crosses were maintained at 29 $^{o}$C. Scale bars: 100 $\mu$m.

### Supplementary Table S1. List of *Drosophila* lines

| *Short name* | *Line Genotype* | *Source or reference* | *Identifier* |
|---|---|---|---|
| ***nub>mRFP*** | w; P{w[nub.PK]=nub-GAL4.K}2, P{w[+mC]=UAS-myr-mRFP}1/CyO | Erkelenz et al., 2021 | originated from RRID:BDSC_63148 |
| ***BuGZ***$^{FlyFos[v318366]}$ | FlyFos016907(pRedFlp-Hgr)(CG1791232384::2xTY1-SGFP-3xFLAG)dFRT | VDRC, Sarov et al., 2016 | |
| ***nub>mRFP; BuGZ***$^{FlyFos[v318366]}$ | w; P{w[nub.PK]=nub-GAL4.K}2, P{w[+mC]=UAS-myr-mRFP}1/CyO; FlyFos016907(pRedFlp-Hgr)(CG1791232384::2xTY1-SGFP-3xFLAG)dFRT | this study | |
| ***BuGZ***$^{RNAi[TRiP.JF02830]}$ | y[1] v[1]; P{y[+t7.7] v[+t1.8]=TRiP.JF02830}attP2 | BDSC | RRID:BDSC_27996 |
| ***BuGZ***$^{RNAi[KK104498]}$ | w; UAS-BuGZ$^{RNAi[KK104498]}$ {30B} | VDRC | |
| ***Lip4***$^{RNAi[TRiP.HM05136]}$ | y[1] v[1]; P{y[+t7.7] v[+t1.8]=TRiP.HM05136}attP2 | BDSC | RRID:BDSC_28925 |
| ***myc***$^{RNAi[TRiP.JF01761]}$ | y[1] v[1]; P{y[+t7.7] v[+t1.8]=TRiP.JF01761}attP2 | BDSC | RRID:BDSC_25783 |
| ***myc***$^{RNAi[TRiP.HMS01538]}$ | y[1] sc[*] v[1] sev[21]; P{y[+t7.7] v[+t1.8]=TRiP.HMS01538}attP2 | BDSC | RRID:BDSC_36123 |
| ***myc***$^{RNAi[GD2948]}$ | w;; UAS-myc$^{RNAi[GD2948]}$ | VDRC | |
| ***Actβ***$^{RNAi[TRiP.JF03276]}$ | y[1] v[1]; P{y[+t7.7] v[+t1.8]=TRiP.JF03276}attP2 | BDSC | RRID:BDSC_29597 |
| ***ftz-f1***$^{RNAi[TRiP.JF02738]}$ | y[1] v[1]; P{y[+t7.7] v[+t1.8]=TRiP.JF02738}attP2 | BDSC | RRID:BDSC_27659 |
| ***yki***$^{RNAi[TRiP.JF03119]}$ | y[1] v[1]; P{y[+t7.7] v[+t1.8]=TRiP.JF03119}attP2/TM6B | BDSC | RRID:BDSC_31965 |
| ***Atf3***$^{RNAi[TRiP.JF02303]}$ | y[1] v[1]; P{y[+t7.7] v[+t1.8]=TRiP.JF02303}attP2 | BDSC | RRID:BDSC_26741 |
| ***DH44***$^{RNAi[TRiP.JF01822]}$ | y[1] v[1]; P{y[+t7.7] v[+t1.8]=TRiP.JF01822}attP2 | BDSC | RRID:BDSC_25804 |
| ***brat***$^{RNAi[TRiP.HM05078]}$ | y[1] v[1]; P{y[+t7.7] v[+t1.8]=TRiP.HM05078}attP2 | BDSC | RRID:BDSC_28590 |
| ***mago***$^{RNAi[TRiP.HM05142]}$ | y[1] v[1]; P{y[+t7.7] v[+t1.8]=TRiP.HM05142}attP2 | BDSC | RRID:BDSC_28931 |
| ***pUWG-RNase H1*** | w;; pUWG-RNase H1/TM6B | this study | |
| ***nub>mRFP; pUWG-RNase H1*** | w; P{w[nub.PK]=nub-GAL4.K}2, P{w[+mC]=UAS-myr-mRFP}1/CyO; pUWG-RNase H1/TM6B | this study | |
| ***pUWG-mCherry*** | w; pUWG-mCherry/CyO, P{ActGFP}JMR1 | this study | |
| ***pUWG***$^{\Delta attB}$***-mCherry*** | w; pUWG$^{\Delta attB}$-mCherry/CyO, P{ActGFP}JMR1 | this study | |

| | | | |
|---|---|---|---|
| ***pUWG-mCherry, nub>*** | w; P{w[nub.PK]=nub-GAL4.K}2, pUWG mCherry/CyO, P{ActGFP}JMR1 | this study | |
| ***pUWG$^{\Delta attB}$-mCherry, nub>*** | w; P{w[nub.PK]=nub-GAL4.K}2, pUWG$^{DattB}$ mCherry/CyO, P{ActGFP}JMR1 | this study | |
| ***SmD3::3xHA$^{[FlyORF\ F003987]}$*** | yw;; pGW SmDS3::3xHA attP 86Fb/TM6B | FlyORF F003987 | |
| ***nub>mRFP; SmD3::3xHA$^{[FlyORF\ F003987]}$*** | w; P{w[nub.PK]=nub-GAL4.K}2, P{w[+mC]=UAS-myr-mRFP}1/CyO; pGW SmDS3::3xHA attP 86Fb/TM6B | Erkelenz et al., 2021 | |
| ***w$^{1118}$*** | w[1118] | BDSC | RRID: BDSC_3605 |

### Supplementary Table S2. List of oligonucleotides and plasmids

**Primers used for cloning**

| *Name* | *Sequence (5'-3')* | *Purpose* |
|---|---|---|
| **mCherry fwd** | TTACTTGTACAGCTCGTCCATG | Cloning of mCherry into the pENTR/D-TOPO |
| **mCherry rev** | CACCATGGTGAGCAAGGGCGAGGA | Cloning of mCherry into the pENTR/D-TOPO |
| **delAttB Frag1 fwd XhoI** | CGCACTCGAGCATTTGTGTGC | Deleting the attB sites from pUWG-*mCherry* |
| **delAttB Frag1 rev** | AAGGGGGCGGCCGCGGTGATGCTGAATTCCTGCAGC | Deleting the attB sites from pUWG-*mCherry* |
| **delAttB Frag2 fwd** | GCAGGAATTCAGCATCACCGCGGCCGCCCCCTTCACC | qPC Deleting the attB sites from pUWG-*mCherry* |
| **delAttB Frag2 rev** | TGCTCACCATGCCATCAGCGGCGCGCCCACCCTTTTACTTG | Deleting the attB sites from pUWG-*mCherry* |
| **delAttB Frag3 fwd** | AGGGTGGGCGCGCCGCTGATGGCATGGTGAGCAAGG | Deleting the attB sites from pUWG-*mCherry* |
| **delAttB Frag3 rev XbaI** | ACGTCTAGACTAGCTTACGTCAATT | Deleting the attB sites from pUWG-*mCherry* |
| **RNase H1- fwd SalI** | ACAAACCATGGGAACCAATTCAGTCGACATGTTACTTCCGCGGTATTTTTGCTGG | Cloning of RNase H1 into the pENTR 4 Dual selection |
| **RNase H1 no TAG rev NotI** | ACAAAGCGGCCGCGAACCATTTTTCTGCTTATACAAGGCGG | Cloning of RNase H1 into the pENTR 4 Dual selection |

**Plasmids**

| *Name* | *Purpose* |
|---|---|
| **pUWG** | Gateway destination expression vector, poly-ubiquitin promotor, C-terminal GFP, Hsp27 terminator |
| **pUWG-*RNase H1*** | Gateway destination vector expressing C-terminally GFP-tagged RNase H1 |
| **pUWG-*mCherry*** | Gateway destination vector expressing mCherry |
| **pUWG$^{\Delta attB}$-*mCherry*** | Gateway destination vector expressing mCherry, attB1 and attB2 sites are missing |
| **pENTR 4 Dual Selection** | Entry vector, Gateway cloning |
| **pENTR/D-TOPO** | Entry vector, Gateway cloning |
